# Label-free assessment of red blood cell storage lesions by deep learning

**DOI:** 10.1101/256180

**Authors:** M. Doan, J. A. Sebastian, R. N. Pinto, C. McQuin, A. Goodman, O. Wolkenhauer, M. J. Parsons, J. P. Acker, P. Rees, H. Hennig, M. C. Kolios, A. E. Carpenter

## Abstract

Blood transfusion is a life-saving clinical procedure. With millions of units needed globally each year, it is a growing concern to improve product quality and recipient outcomes.

Stored red blood cells (RBCs) undergo continuous degradation, leading to structural and biochemical changes. To analyze RBC storage lesions, complex biochemical and biophysical assays are often employed.

We demonstrate that label-free imaging flow cytometry and deep learning can characterize RBC morphologies during 42-day storage, replacing the current practice of manually quantifying a blood smear from stored blood units. Based only on bright field and dark field images, our model achieved 90% accuracy in classifying six different RBC morphologies associated with storage lesions versus human-curated manual examination. A model fitted to the deep learning-extracted features revealed a pattern of morphological changes within the aging blood unit that allowed predicting the expiration date of stored blood using solely morphological assessment.

Deep learning and label-free imaging flow cytometry could therefore be applied to reduce complex laboratory procedures and facilitate robust and objective characterization of blood samples.

## Introduction

### Background

Millions of units of stored red blood cell (RBC) products are transfused each year with an increasing global demand (*1*). RBC transfusions save lives by increasing RBC mass in patients that have low oxygen-carrying capacity due to increased RBC loss (traumatic / surgical hemorrhage), decreased bone marrow production (aplastic anemias), defective hemoglobin (hemoglobinopathies, thalassemias) or decreased RBC survival (hemolytic anemias). Continued technological progress in the preservation and storage of cells has enabled blood banks to store RBCs at 1-6 °C for as long as 8 weeks (*2, 3*). The desired quality of stored blood is to achieve less than 1% haemolysis at the end of storage with an in vivo 24-hour post-transfusion survival rate of greater than 75% (*4, 5*).

During hypothermic storage, RBCs undergo a progressive deterioration in quality that manifests through biochemical and biomechanical changes that adversely influence RBC function. These changes, referred to as “storage lesions”, include metabolic lesions such as declining pH and decreases in glycolytic pathway activity, nicotinamide adenine dinucleotide reducing equivalents (NADH), adenosine 5’-triphosphate (ATP) and 2,3-diphosphoglycerate (2,3-DPG); increasing concentrations of potassium (K^+^), superoxide radical (O_2_^-^) and ferric (Fe^3+^) methemoglobin, causing oxidative damage. Storage lesions also include physiologic lesions such as inefficient O_2_ release and membrane deformation that results in crenations, vesicle budding, membrane loss, and apoptosis of RBCs (*6-10*). These storage lesions are influenced by the length of storage, the blood component manufacturing method, and the characteristics of the blood donor (*11, 12*).

The loss of membrane integrity causes the red cells—which normally exist as smooth biconcave discs (discocytes)—to morph reversibly into an echinocyte (crenated disc-like body), characterized by membrane protrusions or spicula. Eventually, the echinocyte degrades irreversibly into a spheroechinocyte (spherical body) (*6*). An increased presence of spheroechinocytes has been associated with rheological changes and increased viscosity, leading to disturbed capillary blood flow and oxygen delivery (*9*). Therefore, morphology can indicate functional status of RBCs and its degradation may depict the safety and efficacy of the stored blood units.

### Methods to examine morphology of RBCs

Currently, investigation and quantification of these biochemical, physiological, and morphological changes requires complex laboratory procedures, many of which may themselves adversely affect the state of the fragile cells. The standard RBC morphology characterization is done by light microscopy. Here, RBC samples are spread (smeared) on microscopic slides, fixed and stained, and observed under a light microscope. In a typical morphological assessment assay, one hundred cells are randomly chosen for morphological characterization into one of five subclasses: smooth disc, crenated disc, crenated discoid, crenated spheroid and crenated sphere. The RBCs are assigned corresponding shape factors (1.0, 0.8, 0.6, 0.4, and 0.2, respectively), which are then summed to yield a “morphology index” (MI), where higher values indicate higher quality (*13*). The MI is considered a standard metric for quantifying the morphological profile of RBCs during storage. However, this technique is labour intensive, prone to subjective bias, and limited by small sample sizes. Moreover, the smearing may itself affect the state of RBCs, possibly causing unwanted alterations to the sample’s morphological profile, particularly for fragile cells following storage or in diseases such as sickle cell anemia.

In contrast to light microscopy, flow cytometry analyzes cells in fluid suspension and is routinely used in clinical diagnosis and monitoring of hematological disorders (*14*). With a throughput of 50,000 or more cells per second, flow cytometry offers quantitative assessment of large cell populations using signals from emitted fluorescence as well as transmitted and scattered light, with less labour, time, and subjectivity. Due to the lack of spatial information in the data, however, specific fluorescent markers are required to identify particular cell populations, which generally requires multiple rounds of labelling (*14*). This increases expense, decreases throughput, and may itself be damaging to fragile cells.

A more recent cytometric technology, imaging flow cytometry (IFC), incorporates a light microscope objective and spatially registered cameras into a conventional flow cytometry platform to capture images of cells in suspension at high resolution (*15*). IFC captures far more information than conventional flow cytometry, as it can record hundreds of morphological features per imaged channel.

We previously demonstrated the use of an IFC system (ImageStream Mark II, Amnis, EMD Millipore) and its accompanying analysis software, IDEAS, for detecting spheroechinocytes (*16-18*) and characterizing populations of RBCs into different morphological classes. The image gating strategies were largely designed manually, using features related to cell size, shape and intensity patterns. However, subtle changes between certain morphology classes required carefully selecting images from each morphology using the IDEAS *feature finder* tool (*19*). Using Fischer’s discriminant ratio, the tool produced combinations of features that provided the discrimination between each pair of RBC morphology classes, based on individual example cells selected by experts.

To address these issues, we set out to create a less labour intensive and subjective standardised clinical protocol for the assessment of the banked RBCs for transfusion. We hypothesized that machine learning, particularly deep learning, could better delineate RBC morphologies with subtle differences. In previous work, we showed that a remarkable amount of information about cellular phenotype can be captured with few or no biomarkers using brightfield and darkfield images. These features obtained from label-free images are often subtle and invisible to experts yet can be used for classification of phenotypes or cell cycle stage (*20-22*).

Our best results to date were achieved using convolutional neural networks (CNN), a class of machine learning algorithms that has transformed the field of computer vision, including biological imaging. Working directly on pixels in images, rather than on measurements extracted from the entities in the images, CNNs obviate the need for both segmenting objects-of-interest and defining measurements to be extracted from the objects. Designed with hierarchical architectures, CNNs utilize multiple levels of representation, where the representation at one level (starting with the raw imagery input) is subsequently transformed into more abstract representations at higher levels. Higher layers of representation amplify features of the input image that are important for discrimination and suppress irrelevant variations (*23*).

CNNs typically need a large number of images to train the neural network and work particularly well when the object of interest is already isolated (such that no other objects are present in the images). IFC is therefore an ideal match for CNNs, because IFC experiments can produce millions of images of single cells within an hour, each centered in a relatively small frame (~50x50 pixel). The ability of CNNs to automatically generate features—typically in the hundreds to thousands—provides a rich description of cellular morphology, capturing subtle morphological changes that may be unobservable to humans (*22*).

Here, we developed an open-source workflow, called Deepometry (*24*), to facilitate the application of deep learning in cytometry, especially for analyzing IFC data. Deepometry uses a Keras-based implementation of the ResNet50 neural network (*25, 26*), alongside other logistic modules to ease the data flow between raw IFC images and the model. Unlike other deep learning frameworks, which are limited to three-channel RGB images, our modification of ResNet50 allows researchers to use any number of stained or unstained channels.

We present here the application of this workflow to characterize morphological profiles of RBCs at different time points during 42-day storage. We were able discern the differences among multiple RBC morphology classes to ultimately replace manual analysis of blood smears for blood banking.

## Results

### Sample preparation and manual annotation of cell morphologies

To obtain images for deep learning analysis, we analyzed processed samples from whole blood collected from healthy volunteer donors. Samples were processed into red cell concentrates at the Canadian Blood Services (CBS) Centre for Applied Development (netCAD, Vancouver, Canada) in accordance with standard blood bank protocols as described in (*16*), and stored in a hypothermic environment (1-6 °C) throughout the six-week life span (Figure 1A). Three replicate aliquots were sampled from each of 10 blood bags every 3-7 days until expiration. Collected samples were immediately suspended in standard IFC solution and at least 100,000 paired bright field and dark field images of single RBCs were captured per sample by an IFC at high sensitivity. We then delineated six subclasses of RBC morphology—smooth discs, crenated discs, crenated discoids, crenated spheroids, crenated spheres, and other/ambiguous—using gating and the *feature finder* tool in the companion analysis software (IDEAS) (Figure 1B). This delineation is a modified technique of those previously described (*16–18*). The “other/ambiguous” group denoted those disc-like objects that did not share all the typical characteristic features with the other five delineated phenotypes.

**Figure 1.**
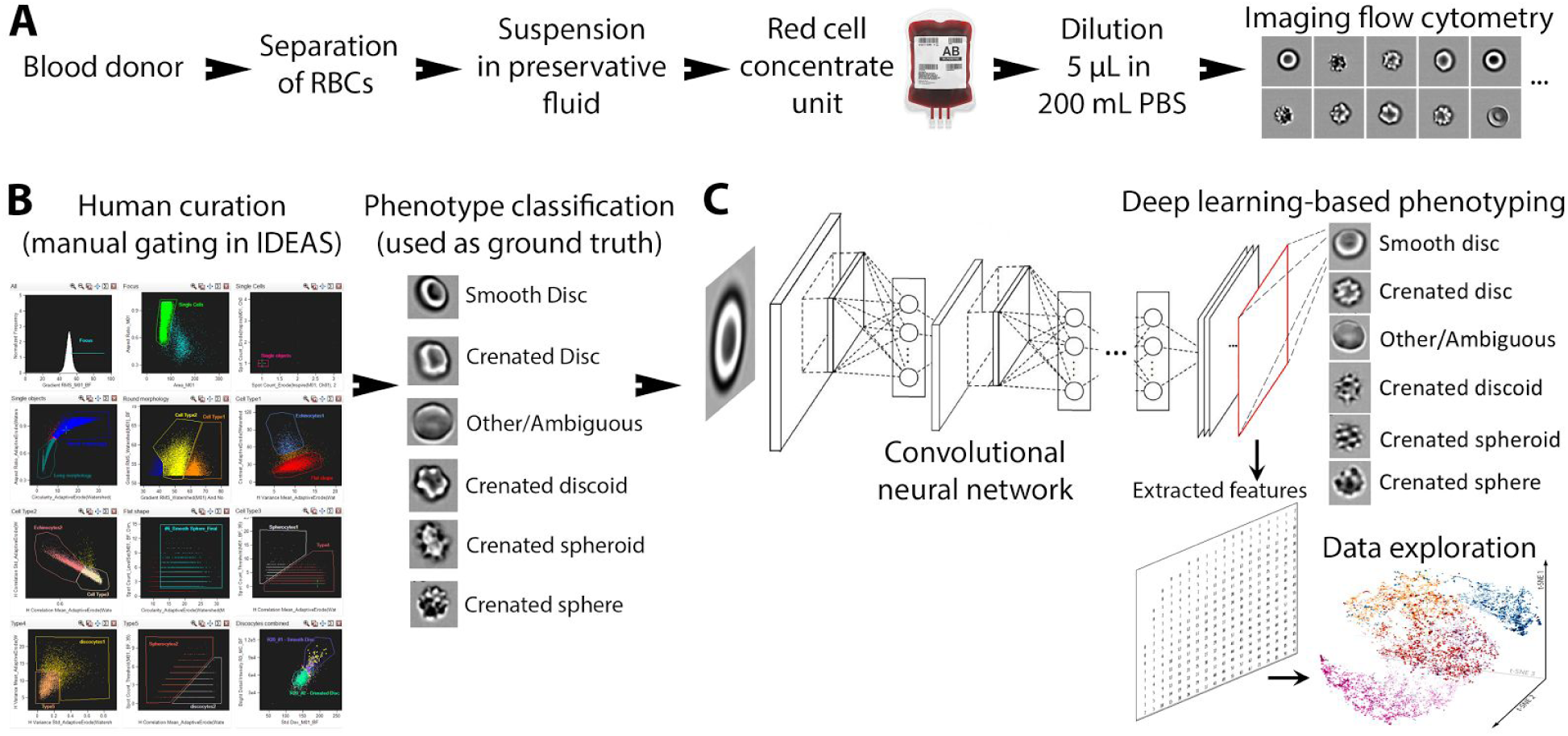
Pipelines for manual / automated assessment of RBC morphology. **A**: Canadian Blood Service staff separated RBCs from donated blood and suspended them in a preservative solution for shipment as a red cell concentrate unit. Samples from these units were then measured by the IFC system to capture bright field and dark field images. **B**: Pipeline for human-curated classification of RBC phenotypes. Using IFC analysis software, IDEAS, 100,000 images of each sample were processed using a human-curated manual gating and filtering template yielding relative populations for six distinct morphologies. The snapshot of the gating template shown is a snippet of a much larger IDEAS template. **C**: Pipeline for automated deep learning, supervised classification and unsupervised learning. The input is an IDEAS file with gates that define various classes. The output is a trained ResNet50 model that can be used to assign a phenotype label for each individual cell in an unseen population (upper right), and to compute morphological features that can be used for data exploration (lower right).

### Deep learning on cell images

We developed the deep learning workflow Deepometry to take as input the exported data from IDEAS, parse it, and pass it to a CNN, ResNet50 (*24*). As shown in Figure 1C and elaborated in the remaining sections of the paper, we then *i*) tasked the trained network to categorize each cell into one of the six expert-defined morphological classes, *ii*) used the network outputs to predict the the age of the blood units, *iii*) performed data-driven clustering and ordering of cells, for validation purposes and to glean additional insights.

To minimize the potential for overfitting (that is, a model that works only for the dataset at hand), the collected data from 10 blood bags—totaling 3,435,401 RBCs—were carefully split into several subsets (Figure 2). First, at each time point, three independently drawn samples were obtained from each bag. For convenience, hereafter a “replicate” represents a series of data at different time points from the same bag. Data from these three replicates were kept distinct from each other in the analysis. The eighteen replicates of six blood bags were used to form the pool of training data.

**Figure 2.**
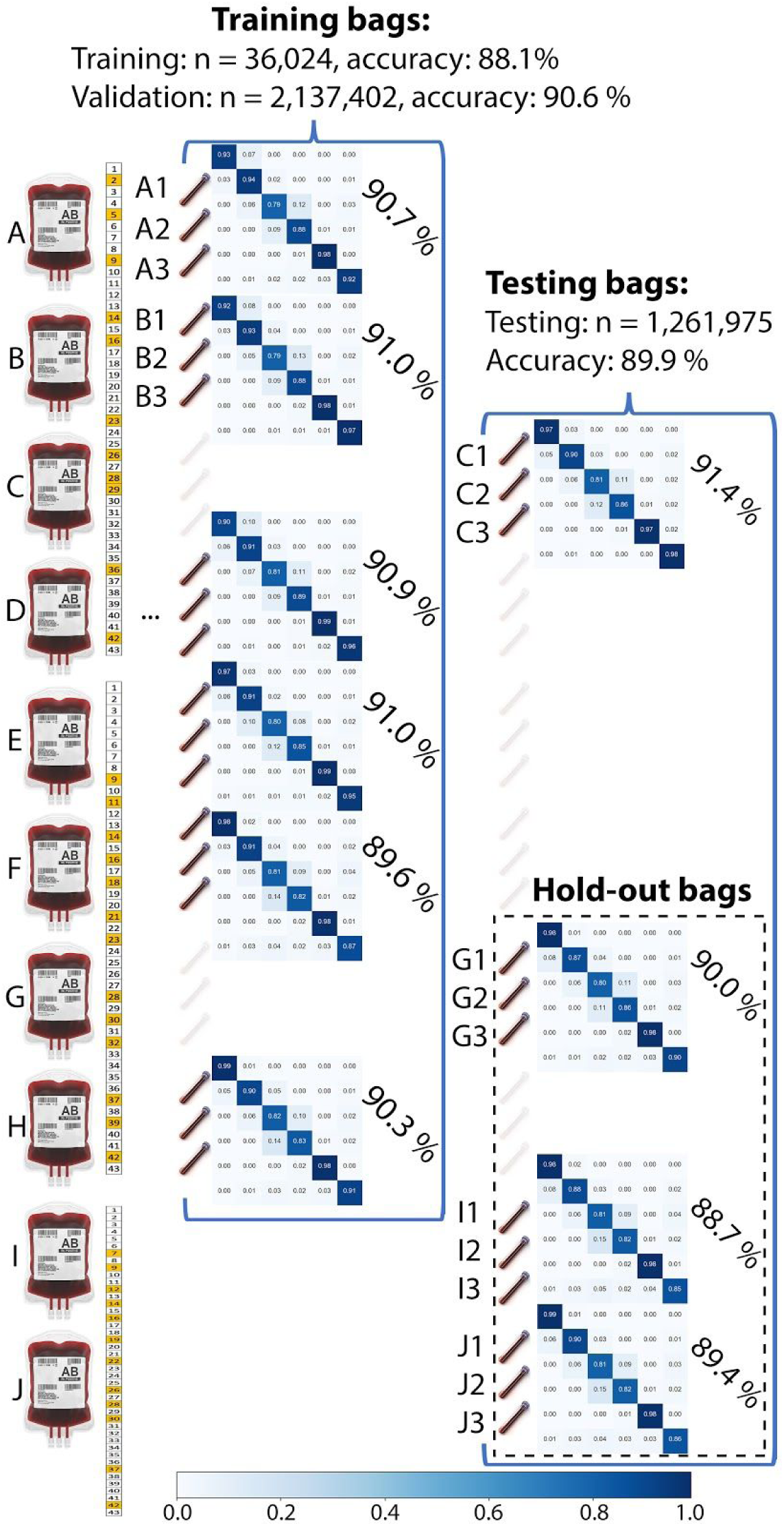
Sampling, training and validating strategies to minimize chances of overfitting. Left: At each time point, each blood unit was sampled and measured three times independently (three replicates per bag). Time points of sample collection are presented on the calendar on the right of each of the three batches of blood bags. The slightly different batches of collecting dates were applied to detect batch effects. Middle: Data from bags A, B, D, E, F and H were pooled together and 1.7% of that pooled dataset was used for training the ResNet50 network. The trained network was then used to perform inference on every replicate of bag A, B, D, E, F and H and prediction on unseen bag C. Retraining then occurred and settings were optimized until the network was considered final. After finalization of the network, the prediction on the previously unused hold-out bags G, I, J was computed a single time. Each confusion matrix shows the prediction of six categories of RBC morphologies performed by the final trained ResNet50. Detailed confusion matrices can be found in Supplementary Figure S1-S3. Weighted prediction recall averaged across all six classes for each blood unit are noted on the right of each confusion matrix.

Only 1.7% (36,024 RBCs) of the cells from these six bags were used for the actual training of the ResNet50 network, including intrinsic validation to optimize the loss and accuracy of the neural network. 98.3% of the pool was then used in an initial prediction test. One blood bag (3 replicates) was kept outside the training pool and was used as an additional (second level) performance test of the trained network. The network was iteratively retrained and improved and the training data refreshed through these two layers of testing until it was declared finalized (Figure 2). The final trained ResNet50 was able to predict the morphological classes of 2,137,402 RBCs in the six pooled training bags with an average prediction recall of 90.6%, as well as 91.4% average recall for the three replicates of the bag used for second level testing.

Three remaining blood bags (9 replicates) were kept held-out and untouched until the final performance test. The trained network was not allowed to be optimized any further during or after this test. Because the held-out data was blocked from any optimization of the study framework, it aims to reflect an unbiased assessment for the study. The finalized ResNet50 network achieved 89.4% accuracy when being challenged by 9 replicates of the held-out samples. Overall, prediction accuracy was 90.3% for 30 tested datasets.

Recall for a given class was consistent across all the replicates of the tested blood bags, as seen in the confusion matrices in Figure 2 and summarized in Figure 3A. The standard deviation of the recall across all samples from 10 bags at various time points (Figure 3A) was less than 1% (0.97%). This indicates that many of the most common sources of variability in such an experiment (overfitting of the model, batch effects based on the sampling day, sampling procedure, and measuring procedure) were either not present or were well-controlled.

**Figure 3.**
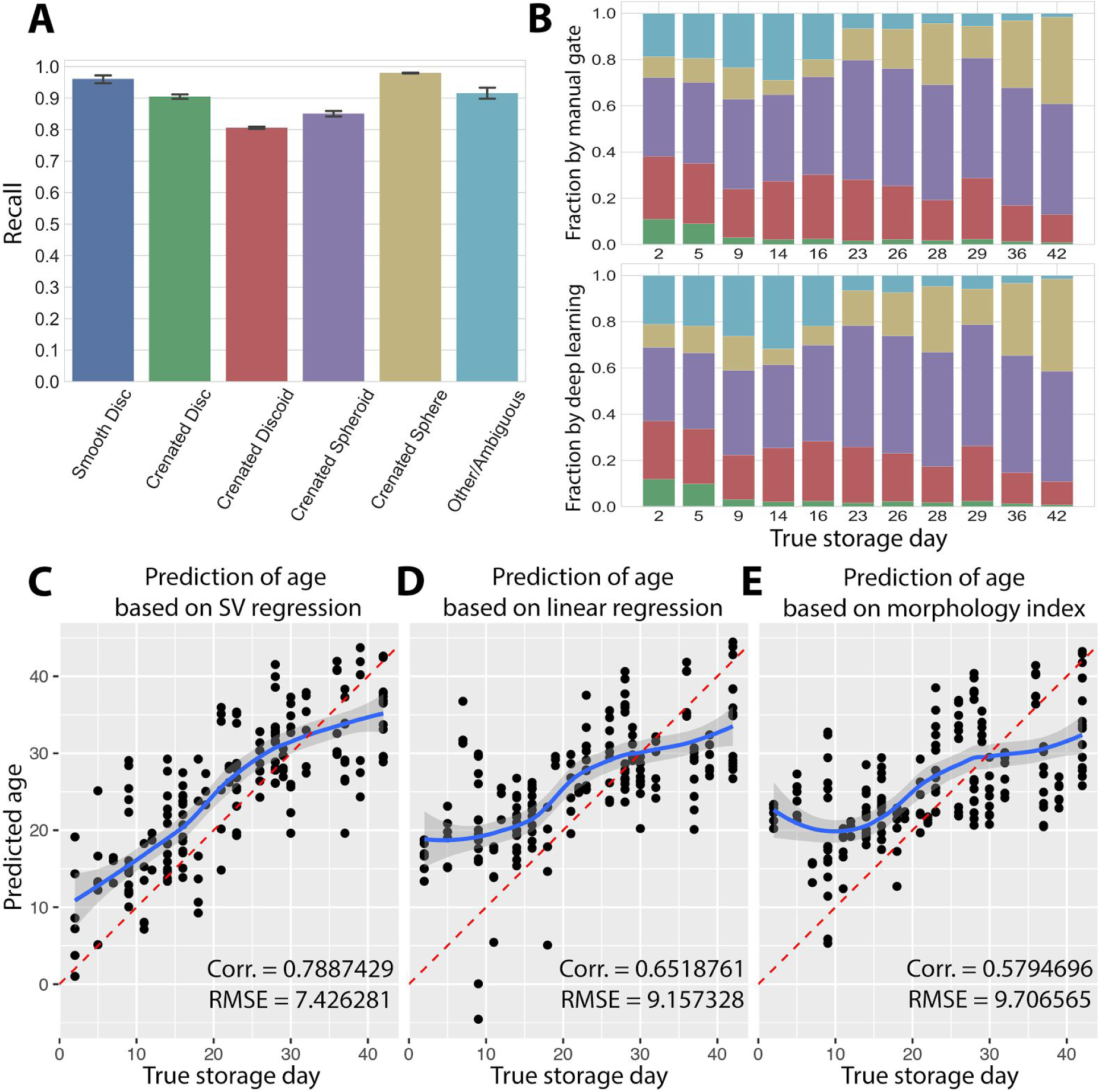
Deep learning model accurately classifies six morphological classes of RBCs and predicts age of blood units. **A**: Prediction recall of a trained ResNet across all replicates and time points of 10 blood bags. Error bars: standard deviation across replicates and timepoints. **B**: Counts for each morphological phenotype and its relative frequency (with respect to total cell number) were calculated at every time point. The frequencies are shown here as stacked fractions. Upper: fractions by human manual gating in IDEAS. Lower: fractions based on deep learning classification. Representative data is displayed from the pooled data of the replicates of four bags (A, B, C, D). Colors of classes are coded in accordance with Figure 3A, but note that the Smooth Disc class is too small to be visible. **C, D**: The direct relationship between the distribution of the six morphologies and the storage age. A support vector regression (C) and a linear regression model (D) were fitted using human manual gating results, which included fraction data from bag C, D, E, F, and J. The fitted models were then used to predict the age of a subset of deep learning results, which included fraction data from bag A, B, G, H, and I. The splitting strategy was aimed to maximize sample variation and minimize batch effects. **E**: Linear regression model of the conventional morphological index. Red lines in C, D, and E show the identity line between true and predicted age. Blue lines are fitted curves of the regression functions. The shaded band is a pointwise 95% confidence interval on the fitted values. Corr.: correlation coefficient, RMSE: root-mean-square error.

Not surprisingly, the prediction recall was strongest (above 90%) for morphologies considered unambiguous by experts, including the smooth disc, crenated disc, crenated sphere (Figure 3A), while that of the transitional morphologies, including the crenated discoid and spheroid, were lower (80-86%).

The distribution of the six morphologies is routinely used by human experts to evaluate the quality of the blood units: the relative frequency of smooth disc, crenated disc, crenated discoid, crenated spheroid and crenated sphere are multiplied with corresponding shape factors 1.0, 0.8, 0.6, 0.4, and 0.2, respectively, which are then summed to yield a “morphology index” (MI), where higher values indicate higher quality (*13*). Mimicking this strategy, we used the automatic categorization of RBC morphologies to count and calculate the relative frequency of each of the six morphologies at each time point. The frequencies (or fractions) calculated by the trained network were nearly identical to that of the manual gating throughout all inspected time points (Figure 3B).

We then examined whether the six fractions of the categorized morphologies could assess the quality of blood bag, and if so, whether it would be statistically comparable to conventional MI, which is a single index. We found that a support vector regression (SVR) model fitted to the six categorized RBC morphologies could accurately predict the storage age of blood bags (Figure 3C). Both linear regression (Figure 3D) and the use of MI (Figure 3E) were less successful at predicting age, especially at early time points.

### Visualization of deep learning feature space and data-driven reordering of RBC morphological progression

RBCs are the most numerous cells in the blood, with a lifespan of more than 100 days. It is quite likely that the current practice of categorizing RBCs into six discrete morphological classes is artificially simplifying a more complex progression between various continuous morphological states, including perhaps even subtle phenotypes overlooked by human experts. A data-driven visualization might reveal unexpected groupings and transition patterns in the natural progression of cell phenotypes. Such an analysis might also serve to validate that the morphological transitions of clinical interest were accurately captured by the deep learning approach, whose features were otherwise relatively inscrutable.

To this end, we used the trained ResNet50 as a feature extractor for the images of all 3.4 million RBCs from 10 blood units across all different time points. The 2048 features generated by the network comprised the high-dimensional representation of each cell. We then used unsupervised dimension-reduction methods such as t-distributed stochastic neighbor embedding (t-SNE) (*27*), diffusion mapping (*28*), and diffusion pseudotime (*29, 30*) to see whether cells would self-organize into meaningful patterns. These methods are complementary; t-SNE generally emphasizes the local dissimilarity between clusters, diffusion map emphasizes the continuum of cell morphology, and diffusion pseudotime highlights the direction of the continuum.

The first two t-SNE components calculated from the extracted deep learning features indeed showed that the six clinically defined morphological types occupy different regions of feature space (Figure 4A). This is somewhat less the case for the first two projected diffusion map components (DCs); as expected for this method, they showed less distinction between different morphological classes (Figure 4B), although the third DC held additional resolving power (not shown).

**Figure 4.**
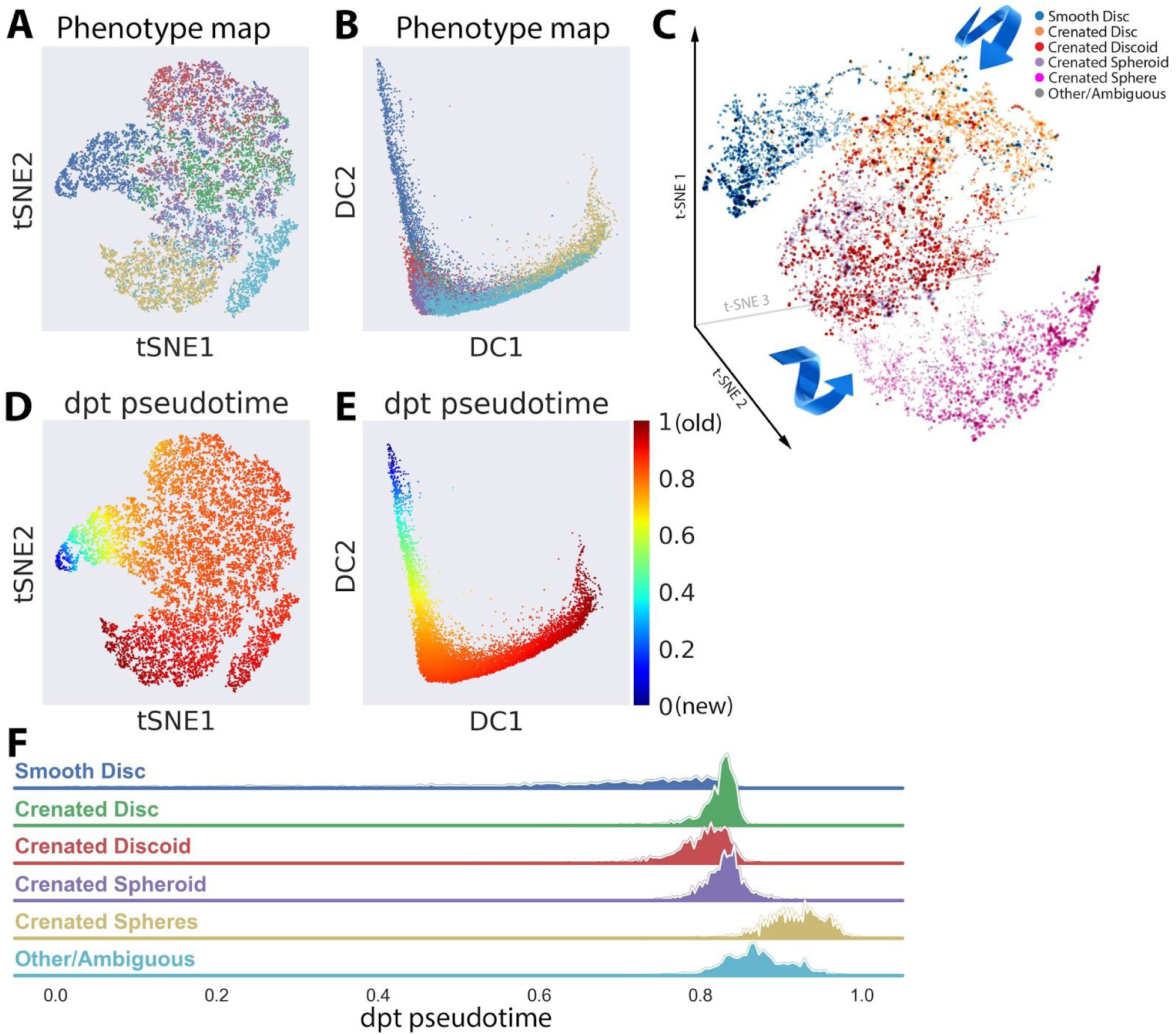
Visualization of deep learning feature space and data-driven reordering of RBC morphological progression. **A-B**: Visualization of deep learning feature space. Shown are RBCs as individual points plotted along the first two components of a t-SNE (A) and diffusion map (B) projection of 2048 deep learning features of the trained ResNet50. Figure 4A, B and F share the same color code, as does Figure 3A-B. Representative data was selected from all replicates of Bag J—one of the hold-out bags—containing 2000 random cells for each class. **C:** Visualization of the first three components of t-SNE calculated from deep learning feature space. The self-ordering of six morphologies was reproducible at 300 t-SNE iteration, perplexity 30 and learning rate 10. Arrows show the speculative direction of transitions between phenotypes. Further 3D inspection is shown in Supplementary S4 movie. **D-E**: The t-SNE and diffusion maps recolored by the cells’ respective pseudotime scores. Figure 4D and 4E are meant to be visualized in pair with Figure 4A and B, respectively. Figure 4D and E share the same color scale. **F**: The prevalence of six phenotypes based on calculated pseudotime score. The Y axis reflects the number of cells in the histogram.

Visualization of the first three t-SNE components in three dimensions further revealed that morphologies that experts consider to be most similar to one another were also positioned in proximity in the 3D tSNE plot (Figure 4C and Supplementary S4 movie). Despite the stochastic nature of t-SNE, an observable transition from one phenotype to another could reproducibly be constructed after about 300 t-SNE iterations (perplexity 30). Moreover, the directional order of morphologies closely resembled the known biology: a smooth disc undergoes a series of intermediate transformations, including crenated disc, crenated discoid and crenated spheroid, before eventually transforming into crenated sphere.

We then assigned diffusion pseudotime values based on diffusion map components and setting a cell that had the highest second-DC value as the initializing root. t-SNE and diffusion map visualizations with overlaid pseudotime values (Figures 4D and E) both confirmed the direction of morphology changes along a continuum, hence the term “pseudotime”. The distribution of the colored clusters in Figures 4D and E were well correlated with the phenotypic clusters in Figures 4A and B, respectively. The Spearman’s rank correlation coefficient ρ (rho) for the relationship between pseudotime and human evaluation of shape factors was −0.59, confirming pseudotime as a measure for the continuous degradation of morphologies through time.

The smooth disc population (blue in Figure 4A, B) was dominant in the early pseudotime (blue in Figure 4D, E), while the crenated sphere population (yellow in Figure 4A, B) was dominant in the late pseudotime (red in Figure 4D, E). The distribution of six phenotypes based on pseudotime are summarized in Figure 4F, reflecting the known biological progression of the morphologies.

Overall, these results lend confidence that the features extracted by deep learning are biologically meaningful, allowing one to gain an unbiased insight that the transition from features describing smooth discs to those of crenated spheres follow the continuum of morphology degradation, further supporting their use in assessment of RBC morphologies.

## Discussion

### Clinical significance of this study

In this work, we demonstrate that deep learning and imaging flow cytometry can be used to evaluate the impact of storage on six clinically-defined RBC morphologies. The proposed technique is well-suited to evaluate other factors known to impact RBC quality, including donor characteristics (age, sex, ethnicity, frequency of donation) and different methods of separating blood components (RBCs, platelets, and plasma) (*11, 12*).

The consistency in scores across bags and replicates indicates that the proposed method may be a reliable tool for evaluating RBC morphological status, although testing across sites is needed to assess consistency across blood component manufacturing methods and from multiple blood manufacturing facilities. Nevertheless, this study provides researchers in the field a useful screening methodology to better understand and predict how donor and manufacturing factors influence RBC storage injury.

If degradation of RBCs follows a strongly consistent and characteristic pattern over time (as suggested in Figure 3C), it may also be possible to derive an RBC morphological profile for individual donors by imaging flow cytometry and deep learning right at the time the blood unit is prepared. This profile can be then used to prospectively predict blood unit quality at various storage time points. In effect, a more precise expiration date could be applied to each blood unit based on its initial state, reducing the need to re-examine the other aging units of the same batch, which voids the sampled bags. Inference of morphological quality across aging units could also allow a blood unit with defined morphology and quality characteristics to be directed to individual patients. This would require a study with a much larger number of blood units from multiple donor groups, preparation procedures, and manufacturing facilities.

Having shown that label-free analysis of RBC morphologies can predict the age of red cell concentrate (RCC) units, we propose that, after further testing, this strategy could replace current manual protocols. The simple sample preparation to be used in this technique would enable clinicians to quickly and easily assess the quality of stored blood, in contrast to microscopic examination (which might adversely smear the sample), conventional flow cytometry (which requires complex laboratory procedures for biomarker labelling), or imaging flow cytometry followed by manual gating (which is time-consuming and subjective).

With proper training on appropriate datasets, the proposed biomarker-free deep learning protocol could also be sufficient to profile various types of blood cells for diagnosis and monitoring of hematological disorders (*31*), as well as in liquid biopsies, for instance, for detection of circulating metastatic/neoplastic cells. We therefore foresee a number of clinical applications being enabled.

### Methodological significance

We developed a robust and reproducible deep learning workflow, publicly disseminated at (*24*), which directly accepts the exported data from IDEAS and uses ResNet50 (*25, 26*) as the deep learning model. The workflows were written in Python, utilized the open-source Keras framework for building the deep learning architecture, and was designed to be user-friendly, requiring minimal expertise in either classical machine learning or deep learning. We believe these robust workflows could be useful for other researchers working with IFC, and therefore disseminated the work via an open-source Github repository (*24*). Furthermore, we facilitated the visualization of deep learning feature space by streamlining the workflow outputs so they can be viewed using public web-based visualization tools, such as http://projector.tensorflow.org and http://clue.io/morpheus/, for interactive inspection.

#### Limitations

The study was conducted with 10 blood bags from a single manufacturing facility, which should be expanded to a larger scale, including a broader range of variety, in term of multiple donor demographics, preparation procedures, and manufacturing facilities.

The prediction power might be improved by further tuning the applied ResNet50, or by using other types of neural network architectures. We however did not aim to deliver a maximally tuned network, which risks becoming specific for this particular task and thus less robust for generalizing to other applications.

It may also yield improvement to train regression models to learn storage age directly from images without the intermediary classification of the six clinically defined morphologies.

#### Steps that need to be taken for the findings to be applied in the clinic

The findings in this study should be validated against conventional biochemical and biophysical assays. RBC morphology characterization could be validated against the clinical gold standard light microscopy technique, using morphology index (MI) as a parameter for comparison as opposed to the gating-based technique used here. If the validation studies show substantial equivalency between the techniques across testing sites, a protocol for an assay involving the IFC system and the deep learning framework for incorporation could be implemented in a clinical setting.

## Materials & Methods

### Sample preparation

Ten red cell concentrate (RCC) units were provided by Canadian Blood Services (CBS) networked centre for applied development (netCAD, Vancouver, Canada). RBCs were separated from anticoagulated donated whole blood using the top/bottom manufacturing method and leukoreduced (*32*).

RCC units were then stored in saline-adenine-dextrose-mannitol at 1-6°C throughout the six-week life span. A sampling site coupler was inserted into each unit for needle aspiration. 18-gauge needles were attached to *safe*PICO syringes (Radiometer, Copenhagen, Denmark) to extract, per unit, three individual 1.5 mL aliquots of RCC. To detect batch effects based on particular collection day or interval between collections, 3 time point collections schemes were used: 11 time points for bags A-D (day 2, 5, 9, 14, 16, 23, 26, 28, 29, 36, 42), 13 time points for bags E-H (day 9, 11, 14, 16, 18, 21, 23, 28, 30, 32, 37, 39, 42), and 12 time points for bags I and J (day 7, 9, 12, 14, 16, 19, 22, 26, 28, 30, 37, 42), respectively. Further details have been described in (*16*).

### Imaging flow cytometry data acquisition and IDEAS analysis

For each extracted sample, 5 μL of RCC was suspended in 200 μL of PBS (magnesium and calcium free) in a 1.5 mL low retention microfuge tube (Sigma T4816-250A) and inserted into an Amnis ImageStreamX Mark II (Amnis, EMD Millipore, Seattle, USA) image flow cytometer (ISx). The ISx was equipped with 488 nm, 405 nm, 561 nm and 642 nm excitation lasers. Bright field illumination was collected in channel 1 (camera 1, 430 nm–488 nm) and dark field signal was collected in channel 12 (camera 2, 560 nm–595 nm). 100,000 pairs of bright field and dark field images of single RBCs were captured using the low speed/high sensitivity settings at 60X magnification. The IFC measurements were repeated every 3-7 days until expiration.

Analysis software IDEAS v6.2 was used to process acquired data using an RBC morphology image gating template as previously described and improved (*16-18*). The first template segregates between four categories (smooth/crenated disc, crenated discoid, crenated spheroid, crenated/smooth sphere). The subclasses that are joined by a “/” implies that the template was unable to further delineate them. A later improved template, which has been used for this work, was able to delineate six morphologies (smooth disc, crenated disc, crenated discoid, crenated spheroid, crenated sphere and other/ambiguous group). The other/ambiguous group denoted those disc-like objects that did not share all the typical characteristic features with the other five delineated phenotypes, and were regarded by human experts as close to crenated disc phenotypes.

The template generates phenotype assignments for each cell as well as absolute counts and relative population percentages for each RBC class.

### Ground truth annotation

In the development of this template, specifically in the selections of masks and features, “truth populations” for each RBC morphology subclass were manually annotated in consultation with an RBC morphology expert. These truth populations are used by the IDEAS *feature finder* tool (*19*) in order to determine the optimal combination of features that best discriminate each RBC morphology class. Images of each unique subpopulation was then exported as a. CIF file.

### Data sampling and splitting strategy

The ResNet50 network was trained with 36,024 RBCs where ~6,000 cells were randomly selected from each of the six classes in pooled data from blood units A, B, D, E, F, H with 3 replicates each (Figure 2). Five out of six cell phenotypes were abundant, however the “Smooth disc” population was rare. This class-imbalance was handled by built-in sampling and augmentation logistic modules; for instance, replicate images of the underrepresented class were generated with various degrees of modification, such as flipping, random rotations, and scaled intensity. The trained network was then tasked to predict the morphological classes of RBCs from the same pooled dataset of 6 bags and 3 replicates of each.

Next, the trained network was challenged by data from bag C. If suboptimal settings were detected during this challenge, a retraining of the fresh network would take place with improved parameters.

Data from blood units G,I,J were kept held-out and untouched during the development and optimization of the study. These held-out data were only unlocked right before submission of the paper for the final verification of the success of the machine learning models.

### Deep learning

Deep learning workflows directly used the exported CIF files from IDEAS as inputs. Bright field and dark field channels were exported. The study was conducted as completely label-free; no fluorescent stains were used in order to avoid concerns about signal spillover between channels. The images were resized to 48x48 pixels by cropping the peripheral background or padding channel-wise with randomly distributed noise sampled from the background. Additionally, cell images were contrast stretched channel-wise to rescale the intensities between the 0.5 and 99.5 percentiles to the full range of uint8, [0, 256).

We implemented ResNet50 (*25*) as the model architecture. We computed categorical cross-entropy as the loss function and accuracy as our metric. The model was compiled using the Adam optimizer with a default learning rate of 0.0001. The learning rate was reduced by a factor of 10 when the validation loss failed to improve for 10 consecutive epochs. Training was set to stop after 50 consecutive epochs of no improvement in the validation loss. Training and validation data was randomly undersampled per blood unit across cell type to create a balanced data set. 70% of sampled data was assigned to the training data set, with the remaining 30% assigned to internal validation of the model during its training.

The data was zero-centered using channel-wise mean subtraction. Means were precomputed from the training set. Mean subtraction and augmentation on training and validation data were performed in real time. Augmentation included random combinations of horizontal or vertical flips, horizontal or vertical shifts (up to 50% of the image size), and rotations up to 180 degrees.

Augmented training and validation data was generated in batches of 256 images to maximize GPU memory resources. We configured the model to train for a maximum of 512 epochs, though early stopping generally terminated training before 200 epochs. Each epoch ran M / 256 steps, with M as number of training samples, to ensure the entire training set was seen once per epoch. Validation occurred once at the end of each epoch, using the entire validation set with validation step K / 256, where K is the number of validation samples.

Prediction metrics included recall, precision, F1-score, and weighted accuracy. Recalls were chosen as our main report, since it is most related to sensitivity and of highest clinical relevance, as well as being the basis of relative frequency calculation for each morphology in a given sample.

### Regression models

A linear regression and a support vector regression (SVR) model were fitted using human manual gating results, which included fraction data from bags C, D, E, F, and J, representing all three collection schemes. The fitted models were then used to test on an unseen subset of deep learning results, which included fraction data from bag A, B, G, H, and I. The 50-50 split was aimed to maximize sample variation and minimize batch effects; whereas the cross-validation (train on manual gating data - test on deep learning data) was aimed to prevent the overfitting of the regression models by inheriting the classification performance of the deep learning network itself. Correlation coefficient and root-mean-square error of the fitting were computed.

### Data visualizations

We extracted the weights and bias of the second-to-last fully connected layer as a filter (highlighted in red in Figure 1C). The set of weights and bias were then applied onto the data to obtain 2048 deep learning features for each object in the dataset. This filtering mechanism was referred to as a “feature extractor” throughout the main text.

Dimension-reduction methods such as T-distributed stochastic neighbor embedding (t-SNE) (*27*), diffusion map (*28*), and diffusion pseudotime (*29, 30*) were then calculated based on those 2048 features and used as embedding components.

Diffusion map organizes data by defining coordinates as dominant eigenvectors of a transition matrix T that describes random walks between data points.

Diffusion pseudotime measure/score is a random-walk-based distance that is based on a sum over weighted Euclidean distances computed using the diffusion map components. For diffusion pseudotime calculation, selection of an initializing object, termed as root cell, was necessary. A cell that had highest second DC was chosen to be the initializing root.

## Acknowledgements

T.R. Turner at Canadian Blood Services is gratefully acknowledged for consultations associated with validating the selection of images for the truth populations used for analysis. T.C. Chang is also gratefully acknowledged for the development of the RBC gating and filtering template on the IDEAS software platform. The Lunenfeld Tanenbaum Research Institute (LTRI) flow cytometry facility is acknowledged for providing access for image flow cytometry experiments. We also gratefully acknowledge the staff of the Canadian Blood Services’ Network for Applied Development (netCAD) and the generosity of the blood donors who made this research possible.

J. Caicedo, M. H. Rohban and S. Singh are gratefully acknowledged for their expert consultations on developing fundamental concepts and critical elements of the machine learning and deep learning frameworks throughout the study.

Funding for this project was provided through a Collaborative Health Research Projects (CHRP) grant (application # 315271 titled “Characterization of blood storage lesions using photoacoustic technologies”), a joint initiative between the Natural Sciences and Engineering Research Council of Canada (NSERC) and the Canadian Institutes of Health Research (CIHR) awarded to the principal investigators M. C. Kolios and J. P. Acker.

A.E.C. and P.R. acknowledge the support of the US National Science Foundation / UK Biotechnology and Biological Sciences Research Council under a joint grant NSF DBI 1458626 and BB/N005163. Funding to P.R. is also provided by the Biotechnology and Biological Sciences Research Council under the grant BB/P026818/1.

There are no conflicts of interest to be declared.

